# Model based safety ranking of opioid drugs using Adversity index

**DOI:** 10.1101/127530

**Authors:** Krishna Asvalayana, Samadhan Ghubade, Sharayu Paranjpe, Anil Gore

**Affiliations:** CYTEL Statistical Software and Services Pvt. Ltd. Pune, India

## Abstract

Annual ADR report counts of opioid drugs are examined to develop a candidate index of overall safety of a drug. Actual counts for various drugs have been sourced from www.vigiaccess.org. A feature found to be common to all drugs considered is that an exponential function adequately describes the pattern of cumulative counts. In the exponential model, the parameter in the exponent (rate constant) is robust and remains the same whether counts are corrected for exposure or not. We propose use of this rate constant as ‘adversity index’ of a drug. Drugs in use can be ranked by value of adversity index, lower value suggesting safer drug.

**Key points:** Cumulative total of annual ADR report counts of opioid drugs follows an exponential pattern. Rate constant in the model is independent of volume of use of the drug. Hence it is a suitable index of overall safety.

## Introduction

Extensive use of opioid drugs is a fact of life today. Economist (April 6, 2017) states that ‘Americans now consume four-fifths of the global supply (of opioid drugs)’. According to American Society of Addiction Medicine, “in 2015, 2 million (Americans) had a substance use disorder involving prescription pain relievers” (http://www.asam.org/docs/default-source/advocacy/opioid-addiction-disease-facts-figures.pdf). Safety aspects of these drugs cannot be overemphasized. According to Centers for Disease Control and Prevention ‘91 Americans die every day from an opioid overdose (https://www.cdc.gov/drugoverdose/epidemic/).

This paper addresses the question of safety comparison amongst opioid drugs. It appears that ‘research comparing opioids to each other in the treatment of people who have chronic pain is quite limited’ and there is ‘little hard evidence about just how the opioids compare to each other in terms of longterm safety and side effects’. (http://consumerhealthchoices.org/wp-content/uploads/2012/08/BBD-Opioids-Full.pdf)

A study comparing the five opioid drugs found that codeine looked much riskier than the other four drugs (hydrocodone, oxycodone, propoxyphene, tramadol) with respect to cardiovascular events and all-cause mortality when they were prescribed for pain not related to cancer [1]. These “…findings on differential risks of various opioids challenge the conventional notion that the safety profiles of opioids are generally interchangeable”. The issue has immediate implications. “If codeine is of middling efficacy for pain and is more risky than other opioids, there would be little reason to use it.” (http://www.health.harvard.edu/blog/the-safety-of-painkillers-20101220915)

It would seem reasonable to suggest that inquiries about safety comparison of opioids are relevant and potentially useful. Approach here is to base the comparisons on counts of spontaneous ADR reports.

ADR reports are a crucial piece in the drug safety jig saw puzzle. These data seem to be underutilized. One common use is in dis-proportionality analysis. Purpose here is detecting a signal related to a specific adverse event of interest. However, it cannot play the role of an index reflecting overall safety level of a drug taking into account all types of adverse events [2]. Also there are reservations about the capacity of this analysis to generate accurate predictions [3]. Instead a model based index may be better suited to provide a bird’s-eye-view assessment of drug safety. What should we expect from such a measure? It should be a single number with the ability to reflect a general trend in ADR report counts. In other words, large value of the measure should go with larger ADR report count and vice versa. Secondly it should be robust and *not* sensitive to volume of exposure. Lastly, it should facilitate safety comparison among drugs even if its absolute value is of limited significance.

There is a formidable difficulty in using a single number for comparison across a group of drugs. The number may depend on annual volume of drug use or exposure. Generally, ADR counts increase as the drug comes into widespread use. That by itself does not mean that the drug is unsafe. Conversely, a drug that is not popular should not be deemed as safer simply because of lower ADR report counts. Instead, to ensure comparability, counts should be normalized/scaled down appropriately. As an example, while comparing two drugs, if one drug is twice as much in use as the other drug, we should divide the ADR count of the first, more popular drug, by two and then compare with the counts in the other drug. Unfortunately, there are multiple ways of normalizing values. As one possibility, you may divide the ADR count by sales figures. Number of prescriptions and number of patients taking the drug are two other alternatives. Which of these should be chosen? Will the index and rank of the drug depend on that choice? FDA document [4] raises these issues. Such difficulties can be eliminated if the index is unaffected by correction for exposure. Fitting a mathematical model to cumulative ADR counts can lead to development of such a suitable index.

## Shape of cumulative ADR report counts and a mathematical model

Consider a graph with year on x axis and cumulative count of ADR reports on y axis. Accumulation helps in ironing out odd values in a year and in generating a smooth behavior. We try to fit an exponential model given by the equation

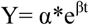

Here Y is the cumulative ADR count at the year t. β is the rate constant. It is easy to show mathematically that β is insensitive to any correction in y for exposure. In other words, if all values of Y are multiplied by 2 and the model fitting exercise repeated, value of α changes but value of β remains the same. This is what makes the exponential model attractive. Of course none of this guarantees that the model will fit the data. Following display shows that the red curve representing cumulative sum of counts for Morphine is followed closely by the fitted line. Notice also that the estimated value of β is 0.145.

Note the following properties of this curve. As value of β increases, the curve rises ever more sharply. If value of β declines the curve becomes flatter. Finally if β reduces to zero, the model becomes a flat horizontal line. It seems therefore that β, the rate constant, is a suitable index to show the pattern of ADR count accumulation.

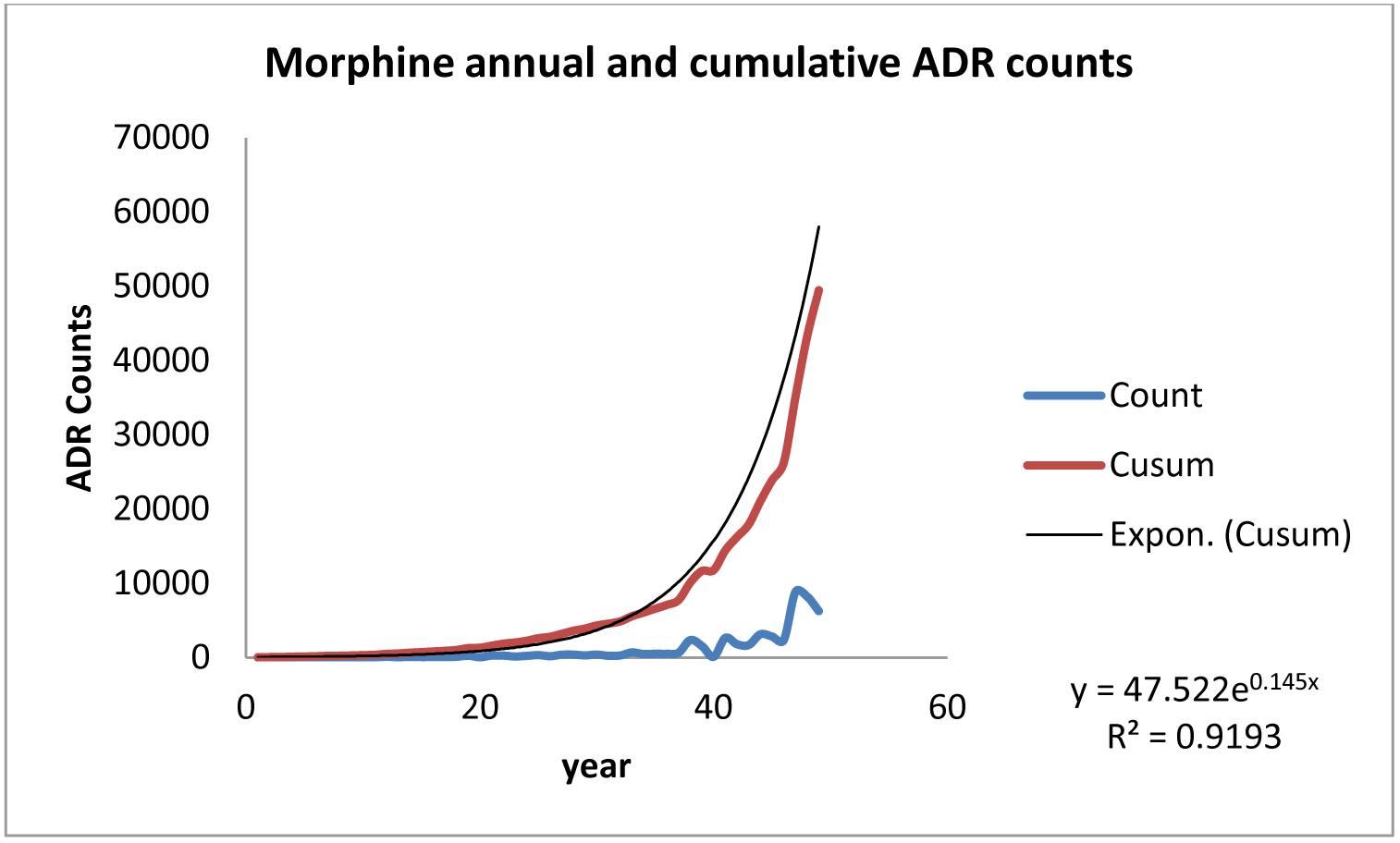

We have considered ten other opioid drugs and repeated the exercise. In all cases, the exponential model is good.

## Ranking of opioid drugs

Following table gives results of fitting the exponential model to a group of 11 opioid drugs. They are arranged in increasing order of β. The smallest value is 0.09 in case of codeine while the largest value is 0.28 in case of Tramadol. Notice that the total ADR count does not follow the ranking. Count for Codeine is NOT the smallest. Count for Percocet is lower.

**Table 1:** Fitting exponential model to cumulative ADR counts and ranking based on growth rate

|  |  |  |  |  |  |
| --- | --- | --- | --- | --- | --- |
| Sr.No. | Drug Name | Adversity index<br>(Growth rate)<br>$\beta$ | $R^2$ | Equation | Total<br>ADR<br>count |
| 1 | Codeine# | 0.09 | 0.83 | $y = 87.366e^{0.0896x}$ | 10487 |
| 2 | Amitriptyline&& | 0.10 | 0.88 | $y = 218.44e^{0.0992x}$ | 22287 |
| 3 | Meperidine (Demerol) | 0.10 | 0.85 | $y = 171.33e^{0.1021x}$ | 22037 |
| 4 | Morphine# | 0.15 | 0.91 | $y = 47.522e^{0.145x}$ | 51387 |
| 5 | Hydromorphone | 0.16 | 0.97 | $y = 3.1465e^{0.1627x}$ | 11662 |
| 6 | Methadone | 0.17 | 0.92 | $y = 5.6807e^{0.1701x}$ | 15349 |
| 7 | Percocet*<br>(acetaminophen +<br>Oxycodone) | 0.20 | 0.95 | $y = 6.2462e^{0.1974x}$ | 8743 |
| 8 | Oxycontin (Oxycodone)# | 0.24 | 0.95 | $y = 0.3156e^{0.2406x}$ | 54966 |
| 9 | Fentanyl | 0.24 | 0.97 | $y = 3.2357e^{0.2407x}$ | 97925 |
| 10 | Hydrocodone (Vicodin) | 0.27 | 0.95 | $y = 3.7189e^{0.2684x}$ | 17601 |
| 11 | Tramadol | 0.28 | 0.93 | $y = 11.893e^{0.2794x}$ | 78850 |

\*short acting #long acting
&& Should this drug be excluded? Does this drug fall in the category in mind?

## Discussion

We have proposed a quantitative method of generating an adversity index for an opioid drug. The index is independent of exposure and hence can be used to compare drugs with different extent of use or popularity. 11 opioid drugs subjected to similar analysis have yielded an overall safety ranking. This should be useful in making a choice in the absence of any patient specific information that points to a subgroup of these drugs. This ranking will have to be validated taking into account other safety information available. Further, it may be of interest to compare counts for a particular organ class instead of the entire body. The approach proposed here can be pursued in such case as well provided the same mathematical model fits data on all drugs under considerations.

Proposed approach to ranking is subject to the condition that all drugs being compared follow the same exponential pattern in the cumulative ADR report counts. In fact this may not always be true. If the common pattern is linear, the method used here will not work. Hence the approach cannot be rigid and fixed. Some flexibility may be needed depending on the case under consideration.

